# Antioxidant activity and phycoremediation ability of four cyanobacterial isolates obtained from stressed aquatic system

**DOI:** 10.1101/311860

**Authors:** Omnia A. M. Badr, Ibrahim I.S. EL-Shawaf, Hoda A. S. El-Garhy, Mahmoud M. A. Moustafa, Omar A. Ahmed-Farid

## Abstract

Cyanobacteria are natural enormous sources of various biologically active compounds with great contributions in different industries. This study aimed to introduce molecular and biochemical characterization for four novel cyanobacterial isolates obtained from Egyptian wastewater canals. Besides, *In vitro* biological activity of these isolates and their potential ability to take up nutrients and heavy metals from wastewater were examined. The obtained accession numbers were KY250420.1, KY321359.1, KY296359.1 and KU373076.1 for *Nostoc calcicola, Leptolyngbya sp, Nostoc sp, and Nostoc sp,* respectively. The isolate *Leptolyngbya sp* (KY321359.1) showed the lowest identity (90%) with other deposited sequences in database. While the isolate *Nostoc sp* (KU373076.1) showed the highest total phenolic content as well as the highest levels of caffeic, ferulic and gallic acids. Consequently, it appeared the highest antioxidant scavenging activity. All cyanobacterial isolates revealed potent ability to take up nutrients and heavy metals from wastewater. Generally, this study provides a taxonomic and molecular evidence for four novel cyanobacterial isolates with antioxidant activity and potent phycoremediation ability.

## Introduction

The continuous increase in populations, industrialization and excessive generation of wastewater are major environmental issues threat sustainability of developing countries. The release of polluted water without appropriate treatment into freshwater resources has led to maximizing the problem of water pollution, besides making the water unfit for drinking, irrigation and aquatic life (Egun 2010). Therefore, there is an urgent need to develop eco-friendly and economic technologies for wastewater treatment, which would require simple infrastructure, lesser inputs and with potential acceptance at the commercial level (Sood et al., 2015).

The ecological and economic importance of cyanobacteria is growing rapidly worldwide due to the great diversity of the products that can be developed from cyanobacterial biomass (Hamed, 2016). The wide range of cyanobacteria biochemical products and the potential use of these compounds in the food, feed, pharmaceutical, nutraceutical, cosmetic and research industries have led to more concern of cyanobacteria (Pulz and Gross, 2004, Griffiths et al., 2016). Commercially, cyanobacteria are used to produce many relatively high-value products such as Carotenoids, β-carotene, astaxanthin and long-chain polyunsaturated fatty acids for use as human nutritional supplements (Borowitzka, 1995, Borowitzka, 2013).

Phycoremediation is broadly defined as the utilization of microalgae and cyanobacteria (blue-green algae), for the removal of contaminants from wastewater. It is a promising technology offer an inexpensive alternative to conventional forms of tertiary wastewater treatments (Sood et. al., 2015). The use of cyanobacteria in wastewater treatment is an eco-friendly process with no secondary pollution as long as the produced biomass allows efficient nutrient recycling (Munoz and Guieysse 2008). Phycoremediation based cyanobacteria is also cost effective as compared to other physical and chemical remediation methods (Han et al. 2007). The high requirement of N and P for the growth of cyanobacteria strengthens their ability to use the nutrient rich wastewater as a medium for multiplication of these organisms. At the same time, assimilated nitrogen and phosphorus can be recycled into algal biomass as biofertilizer (Pittman et al. 2011).

Many beneficial cyanobacterial strains belong to the order Nostocales. The importance of these strains refers to their capacity to improve quality and fertility of soils and retain water. They have the ability to release phosphate, nitrogen, trace elements, decrease chemical nitrogen demands. Also, they produce substances with antiviral and antibacterial activities (Hamed, 2016). Additionally, Nostocales have the ability to take up nutrients from water offer the feasibility to recycle the nutrients into algae biomass and thus can be used in wastewater treatment (Becker, 1994, Sood et al., 2015).

For these findings, cyanobacteria have become one of the most serious tracks for emerging problems that are encountered nowadays (Stephens et al., 2013, Hamed, 2016). Despite their many advantages, of the 50000-existent species, only a few thousand are now kept in collections and are investigated for their chemical content, and even fewer are cultivated in industrial quantities. Therefore, cyanobacteria is still not a well-studied life form from a biotechnological point of view (Mishra et al., 2015; Hamed, 2016).

Therefore, the present study aimed to identify and characterize four cyanobacterial isolates obtained from Egyptian irrigated drainage canals. Molecular characterization based on 16s rDNA, morphological and biochemical characteristics were determined. Moreover, some biotechnological applications of the obtained isolates have been introduced through investigating the *in vitro* biological activity of these isolates and, their potential ability to uptake nutrients and heavy metals from the used wastewater.

## Materials and Methods

### Samples collection

The study was performed in Biotechnology Lab II, Genetics and Genetic Engineering Dept., Faculty of Agriculture, Benha University, Egypt. The samples were collected from irrigated drainage canals, qalyubia governorate. Water samples were collected in sterile plastic bottles and analyzed within 4 h after their collection under sterilized conditions.

### Purification and cultivation of cyanobacterial isolates

Cyanobacterial isolates were purified with serial dilutions according to Lee et al. (2014). They were isolated and purified using nutrient selective media optimized for the growth of cyanobacterial species (BG-11 and modified BG-11).The modified BG-11 medium was prepared from the BG-11 recipe by removing all nitrogen forms to support only heterocystous cyanobacteria (Allen, 1968; Rippka et al., 1979).

### DNA extraction and PCR amplification

Total genomic DNA from pure isolates was extracted according to Fawley and Fawley, (2004). The DNA was amplified using universal 16S rRNA primers 27F:5’. AGAGTTTGGATCMTGGCTCAG.3’and1492R:5’CGGTTACCTTGTTACGACTT. 3′. The used primers were tested by *in silico* PCR tool (http://insilico.ehu.eus/PCR/). The expected PCR amplicon was almost 1.5 kb. PCR reaction was performed in a 50 μl mixture containing 0.4 μM of each primer with concentration of 10 pM, 400 μM of dNTPs mix, 5 μL of 10x PCR reaction buffer, 2 μM MgCl_2_, 2.5 units of TAKARA Taq DNA polymerase (Cat. #: R001AM), 1 μL of template DNA and the final volume was adjusted with sterilized double distilled water. PCR thermocycler (AriaMx) was used to amplify the reactions consisting of 95^0^ C for 3 min followed by 35 cycles at 95^0^ C for 50 secs, 55^0^ C as annealing temperature for 1 min with an extension of 72^0^ C for 1 min followed by final extension temperature at 72^0^ C for 10 min. Amplified PCR products were stored at -20^0^C for further purification and downstream application, then 5 μl of PCR amplified product was loaded on 1.2% agarose gel electrophoresis stained with Ethidium bromide using GeneRuler™ 1kb DNA ladder (Cat. #: SM0313), then visualized under UV Transilluminator (Bio RAD).

### Cloning and Sequencing

The expected DNA band,1500 bp, was eluted from agarose gel and purified according to the manufacturer’s QIAquick Gel Extraction Kit (Cat. #: 28704). The purified PCR fragment was ligated into pGEM^R^-T Easy Vector Systems (Cat. #: A1360) according to the manufacturer. The competent cells of *E. coli* top 10 strain were prepared and transformed as described by Inoue et al. (1990). The white colonies were picked up from LB/Amp/Xgal plates and inoculated on LB/Amp broth media, then incubated overnight at 33 °C with shaking to stabilize the plasmid inside the transformed cells. The alkaline method of Bimboim and Doly (1979) was used to isolate the plasmid. The purified plasmids were analyzed using electrophoresis on 1.2% agarose gel using GeneRuler™ 1 kb DNA Ladder to confirm the recombinant plasmids which were sequenced by Macrogen Company, South Korea.

The obtained sequences for 16S rRNA gene were examined for vector contamination using the VecScreen tool (http://www.ncbi.nlm.nih.gov/tools/vecscreen).While Jalview software (Waterhouse et al., 2009) was used to show single nucleotide polymorphisms (SNPs) and consensus resulted from the alignment of our obtained sequence and the nearest strains in NCBI database (http://www.jalview.org/). Construction of the phylogenetic tree was done using Clustal Omega (https://www.ebi.ac.uk/Tools/msa/clustalo/) and MEGA7 software (Kumar et al., 2016). The obtained sequences from irrigated drainage water canals were registered at NCBI database under accession numbers KY250420.1, KY321359.1, KY296359.1 and KU373076.1 for *Nostoc calcicola, Leptolyngbya sp, Nostoc sp, and Nostoc sp,* respectively, (http://www.ncbi.nlm.nih.gov).

### *In vitro* biological activity of cyanobacterial isolates

Determination of flavonoids, vitamin A and E, water-soluble vitamins, β-carotene and Zeaxanthin were determined by HPLC (Nogata et al., 1994; Gimeno et al.,2000; Ekinci and Kadakal, 2005 and Ahamed et al., 2007). The total phenolic content of the extracts was determined by the Folin-Ciocalteu method (Kaur and Kapoor, 2002). The ability of different extracts to act as hydrogen donors was measured by 1, 1-diphenyl-2-picrylhydrazyl (DPPH^) assay as described by Blois (1958).

### Phycoremediation ability of cyanobacterial isolates

The identified cyanobacterial isolates were cultivated on autoclaved wastewater media obtained from Moshtohor sewage canal under continuous lighting (1000 lux) as a batch culture in 250 ml Erlenmeyer flasks containing 100 ml of BG-11 at 27 ± 2 °C. To determine the ability of these isolates to uptake nutrients and heavy metals from wastewater. The nutrients and heavy metals concentration were measured by Perkin EL Mer 3300 Atomic Absorption Spectroscopy before cyanobacteria cultivation and at the end of the experiment. The uptake of heavy metals by the cyanobacteria was calculated by the difference of heavy metals and nutrients as percentage removal; (B-A)/B X 100 where, B; control concentration of nutrients and heavy metals in wastewater media before cyanobacteria cultivation and A; concentration of nutrients and heavy metals in wastewater media after cyanobacteria cultivation period. Wastewater medium was filtrated through Whatman filter paper (No. 42) before final measurement. The experiment lasted for 16 days assuming all the heavy metals removed from the water were taken up by the grown cyanobacteria.

### Statistical analysis

The values were expressed as a mean ± SE for 5 replicates of different cyanobacterial isolates. Means and standard error were calculated using SAS software for Windows, version 13.0, (SAS, 2004).

## Results and Discussion

Cyanobacteria represent an extremely diverse group of prokaryotic organisms found in a wide range of environmental systems (Hu et al., 2008). The number of cyanobacteria species ranges from 2,000 to 8,000 but there are only 2,698 described species of cyanobacteria (Nabout et al., 2013). Therefore, little is known about cyanobacteria biodiversity and the number of species in the world (Guiry, 2012).

Qalyubia irrigated drainage canals characterized with the presence of various wastewater types, human sewage, livestock wastes, agro-industrial wastes, industrial wastes, piggery effluent, food processing waste and other agricultural waste substrates. Many investigations have been conducted the impact of some environmental stresses on the distribution and species composition of fresh water algal communities (Abdel-Raouf et al. 2012). Hence, there are a potentiality to find new algal species in such aquatic stressed system. In this study, the first microscopic investigation confirmed that there were many different shapes of diatoms and cyanobacteria and the variation in the obtained isolates may be due the presence of various wastewater types in qalyubia irrigated drainage canals. Serial dilutions method resulted in purifying four cyanobacterial cultures. Three isolates were belonged to *Nostocaceae* family and one belonged to the family of *Pseudanabaenaceae*.

### Molecular characterization of the cyanobacterial isolates

The amplified PCR products of *16S* rRNA genes almost 1.5kb (Fig 1) from cyanobacterial isolates were sequenced. The assembled sequences were deposited in NCBI database under accession numbers KY250420.1, KY321359.1, KY296359.1 and KU373076.1 for *Nostoc calcicola, Leptolyngbya sp, Nostoc sp, and Nostoc sp,* respectively. Analysis of the obtained sequences via VecScreen tool showed no contamination with vector sequence. While the BLAST alignment showed that the nearest deposited sequences in database to the obtained sequences were *Nostoc calcicola* 99 (GQ167549.1), *Leptolyngbya sp. NIES-504* (LC319755.1), *Nostoc sp.* PAN-549 (KF921498.1) and *Nostoc sp. UIC 10279* (JX962720.1) with identity ratio 99%, 90%, 98% and 99% as shown in Fig. 2A, B and Fig.3A, B, respectively.

**Figure 1.**
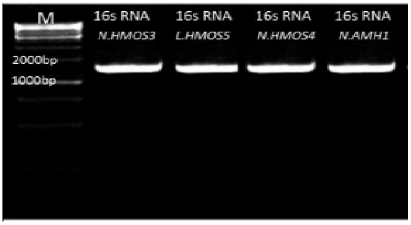
PCR products of 16S rRNA gene were almost 1.5 Kb of *Nostoc calcicola* (KY250420.1), *Leptolyngbya sp* (KY321359.1), *Nostoc sp* (KY296359.1), *Nostoc sp* (KU373076.1) with 1Kp DNA ladder.

**Figure 2.**
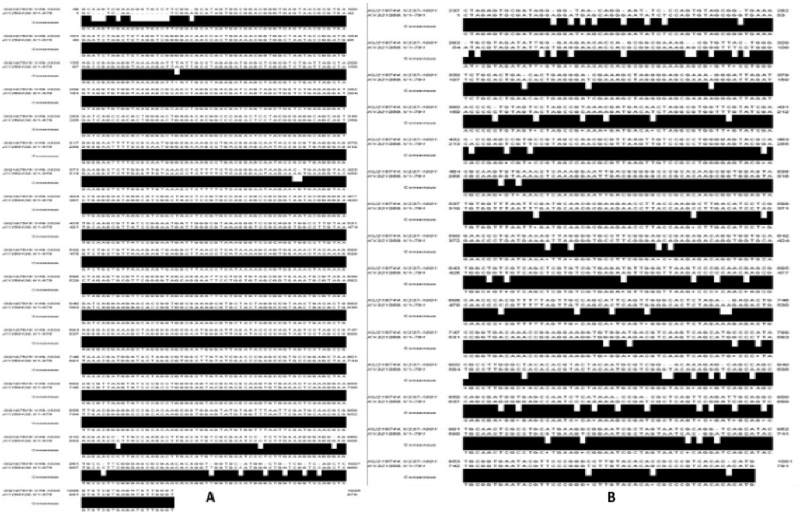
**A**. Image showed many SNPs between *Nostoc calcicola* (KY250420.1) and the nearest one *Nostoc calcicola (GQ167549.1)* deposited in GenBank Database. **B**. Image showed many SNPs between our obtained sequence of *Leptolyngbya sp* (KY321359.1) and the nearest one *Leptolyngbya sp.* (KU219744.1) deposited in GenBank Database.

**Figure 3.**
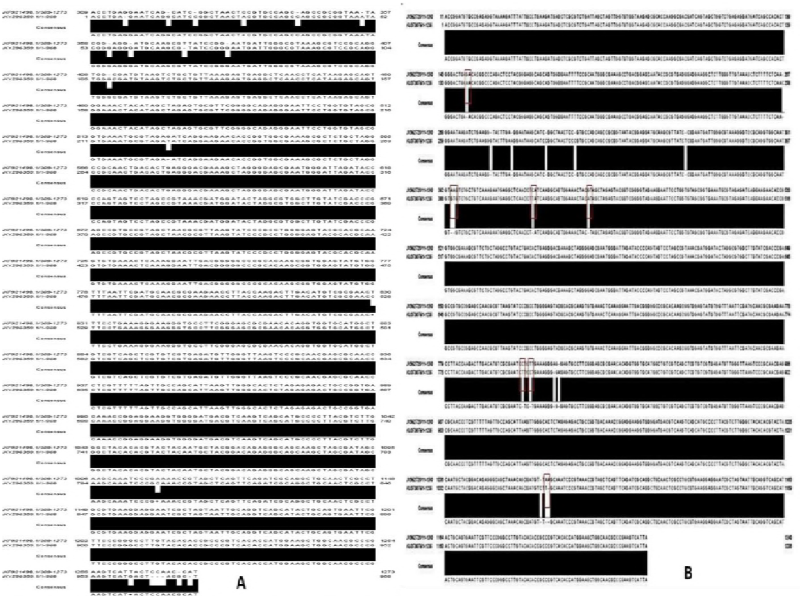
**A**. Image showed many SNPs between *Nostoc sp* (KY296359.1) and the nearest one *Nostoc sp. PAN-549* (KF921498.1) deposited in GenBank Database. **B**. Image showed many SNPs between our obtained sequence of *Nostoc sp* (KU373076.1) and the nearest one *Nostoc sp. UIC 10279* (JX962720.1) deposited in GenBank Database.

The isolate *Leptolyngbya sp* showed the lowest identity ratio (90%) with other deposited sequences in database where, the divergence seemed in high number of SNPs (54) and GAPs (26) between it and the nearest sequence. These results supported by Thompson et al. (2013) who reported that strains from different microbial species share less than 95% Average Nucleotide Identity (ANI). Hence, this isolate could be a new species with novel and unique characteristics and this could be attributed to the area of isolation where high concentration of pollutants is found in the irrigated wastewater cannels. The phylogenetic tree confirmed the same identity ratios on the roots of clades. It clearly diverged from the nearest sequence *(Leptolyngbya sp. NIES-504,* LC319755.1,) and found to be developed in a new clade sharing the same ancestor with the family of Pseudanabaenaceae (Fig 4B). In contrast, the obtained isolates *Nostoc calcicola, Nostoc sp* (KY296359.1), *and Nostoc sp* (KU373076.1) were in the same clades with the nearest sequences *Nostoc calcicola* 99 (GQ167549.1), *Nostoc sp.* PAN-549 (KF921498.1) and *Nostoc sp. UIC 10279* (JX962720.1) in database (Fig 4A and Fig. 5A, B). These results reflect close similarity that seemed in the less number of SNPs and GAPs among the obtained sequences and the nearest sequences in database. The expected restriction maps of all obtained sequences (Fig 6) displayed different endonucleases sites that could be important in building genetic maps and biodiversity studies.

**Figure 4.**
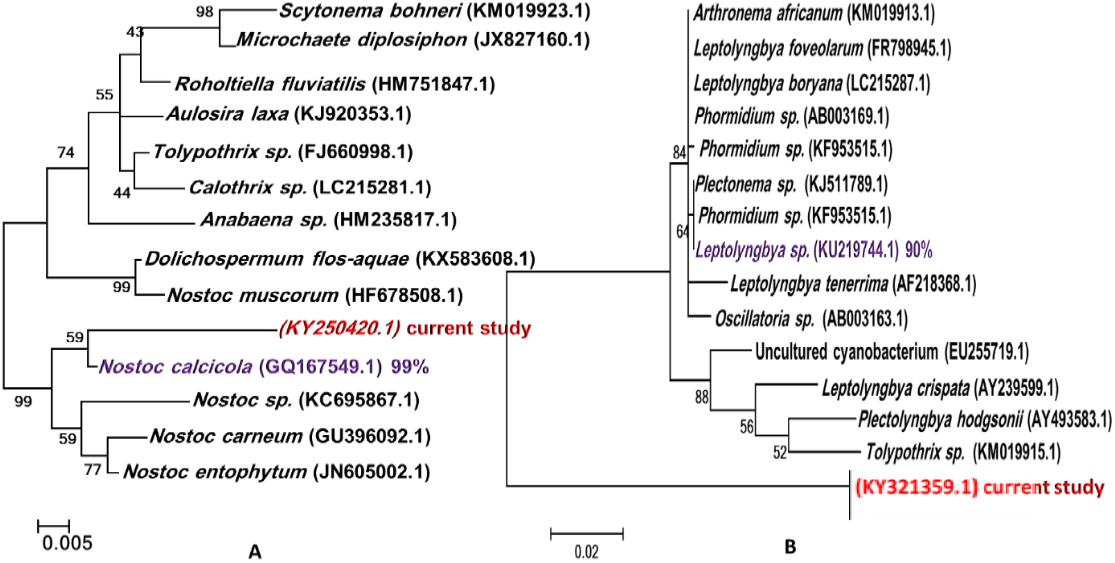
**A**. The phylogenetic tree of the obtained sequence *Nostoc calcicola* (KY250420.1) with the nearest one *Nostoc calcicola (GQ167549.1)* **B**. The phylogenetic tree of the obtained sequence of *Leptolyngbya sp* (KY321359.1) with the nearest deposited sequence *Leptolyngbya sp.* (KU219744.1) in GenBank. The phylogenetic trees were recovered by Maximum Likelihood method using MEGA 7.0.2.1 software. Average Bootstrap values, of compared algorithms, were indicated at the branch roots. The bar was represented 0.005 and 0.02 changes per nucleotide. Accession numbers of database extracted sequences were in black, Purple and red colors for the nearest and the obtained sequences, respectively.

**Figure 5.**
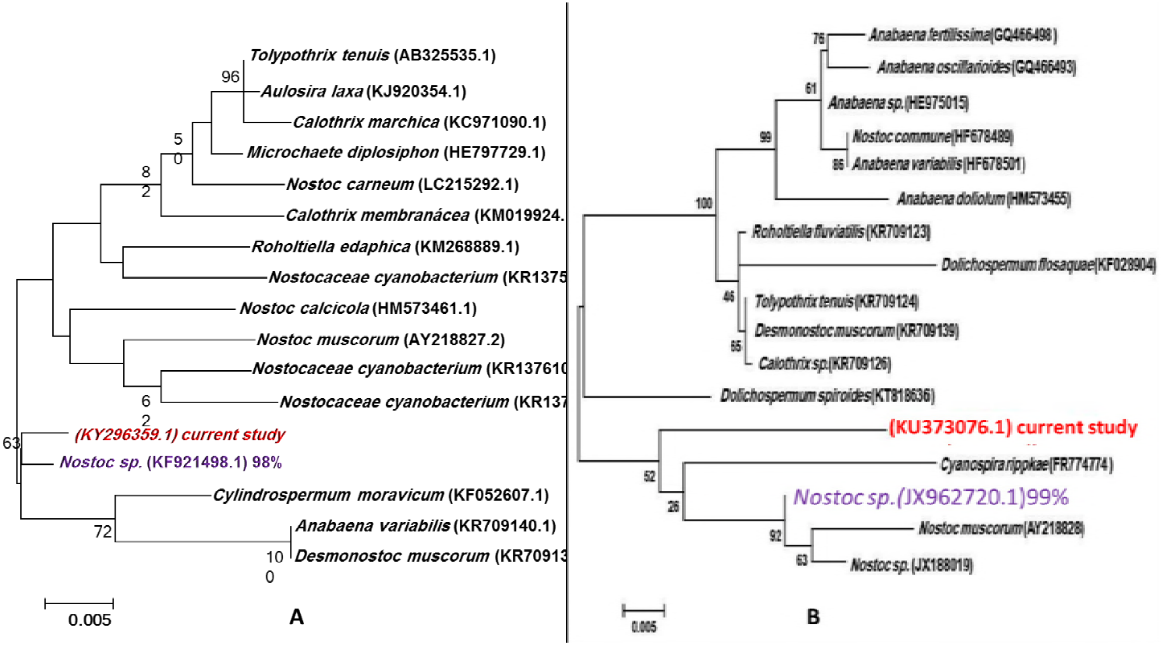
**A**. The phylogenetic tree of the obtained sequence *Nostoc sp* (KY296359.1) with the nearest one *Nostoc sp. PAN-549* (KF921498.1) **B**. The phylogenetic tree of the obtained sequence of *Nostoc sp* (KU373076.1) with the nearest deposited sequence *Nostoc sp. UIC 10279* (JX962720.1) in GenBank. The phylogenetic trees were recovered by Maximum Likelihood method using MEGA 7.0.2.1 software. Average Bootstrap values, of compared algorithms, were indicated at the branch roots. The bars were represented 0.005 changes per nucleotide. Accession numbers of database extracted sequences were in black, Purple and red colors for the nearest and the obtained sequences, respectively.

**Figure 6.**
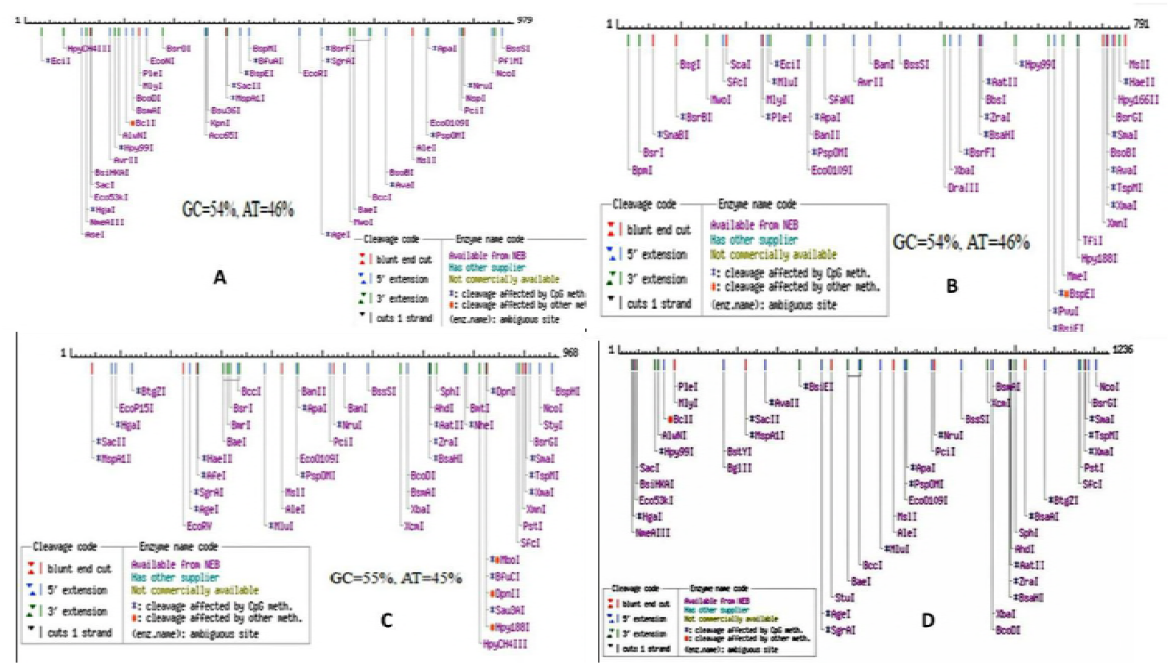
Restriction map of the partial sequences of 16S rRNA genes for A-*Nostoc calcicola* (KY250420.1) *B-Leptolyngbya sp* (KY321359.1) *C-Nostoc sp* (KY296359.1) *D-Nostoc sp* (KU373076.1) with available commercially restriction enzymes

### Biochemical profile of the cyanobacterial isolates

The recent study of Singh et al. (2017) clearly demonstrated that cyanobacteria contain a wide range of carotenoids, flavonoids, and phenolic compounds with a potential Antioxidant, Anti-Inflammatory, Anticancer, Anti-Diabetes, and Antibacterial Activities. Phenolic compounds are considered as one of the most important classes of natural antioxidants (Machu et al., 2015). Some investigations reported that microalgae and cyanobacteria are natural sources of phenolic compounds but, few studies have focused on their identification and quantification in microalgae and cyanobacteria (Cirulis et al.,2013; Safafar et al.,2015; Jerez-Martel et al.,2017). Jerez-Martel et al. (2017) reported that Phenolic constituents were not detected in *Nostoc sp., Leptolyngbya protospira, Nodularia spumigena, Phormidiochaete sp., Arthrospira platensis,* and the strain *Caespitella pascheri* as well as he reported that among the studied cyanobacterial isolates gallic acid was only identified in *Nostoc commune.* However, **Klejdus et al. (2009)** stated that *Spongiochloris spongiosa, Spirulina platensis, Anabaena doliolum, Nostoc sp., and Cylindrospermum sp.* seemed phenolic compounds at μg levels per gram of biomass. The results here agree with the current study results, where phenolic compounds detected in all cyanobacterial isolates and the phenolic content varied from 13.85 in *Nostoc sp* (KU373076.1) to 6.25 in *Nostoc calcicola* (KY250420.1) μg gallic/g dried cyanobacteria Table.1. Moreover, the Nutritional profile of *Nostoc sp* (KU373076.1) recorded the highest levels of Ferulic acid, Coumaric acid, Cinnamic acid, Vitamin A, Riboflavin, Pyridoxine, Zeaxanthin, Gallic and Salycilic acid Table.2. Besides, the caffeic acid levels were high against the results that obtained by Goiris et al. (2014) who screened flavonoids in different microalgal and cyanobacterial species and reported that none of the algal biomass samples contained the flavanone eriodictyol (nor its precursor caffeic acid). Hence, *Nostoc sp* (KU373076.1) is considered very promising as natural antioxidant.

The DPPH (2,2-diphenyl-1-picrylhydrazyl) radical scavenging assay is used widely for evaluating natural antioxidant due to its stability, simplicity and reproducibility (Kuda et al., 2007). The ability of a compound to scavenge DPPH radicals is dependent on their ability to pair with the unpaired electron of a radical; the higher the DPPH-scavenging activity, the higher the antioxidant activity of the sample (PARK et al., 2004). From the results presented in Table 3, all cyanobacterial isolates exhibited high values of capacity to scavenge free radical DPPH that ranged from 18.25% *(Nostoc calcicola,* KY250420.1,) to 29.33% (*Nostoc sp,* KU373076.1,) at concentration of 50 μg/ml. In the current study, Cyanobacterial isolates displayed considerably stronger relative radical scavenging efficiencies in compare with the study of Jerez-Martel et al. (2017) who determined the antioxidant activity of *Nostoc sp., Leptolyngbya protospira, Nodularia spumigena, and Phormidiochaete sp,* where DPPH radical ranged from 7.65% *(Leptolyngbya protospira)* to 27.89% *(Nostoc sp.).* The DPPH scavenging ability is a significant indicator of potential antioxidant activity that may be due to the presence of phenolic compounds and flavonoids in the crude extracts of the isolates as also indicated by Hossain et al. (2016).

**Table 1.**
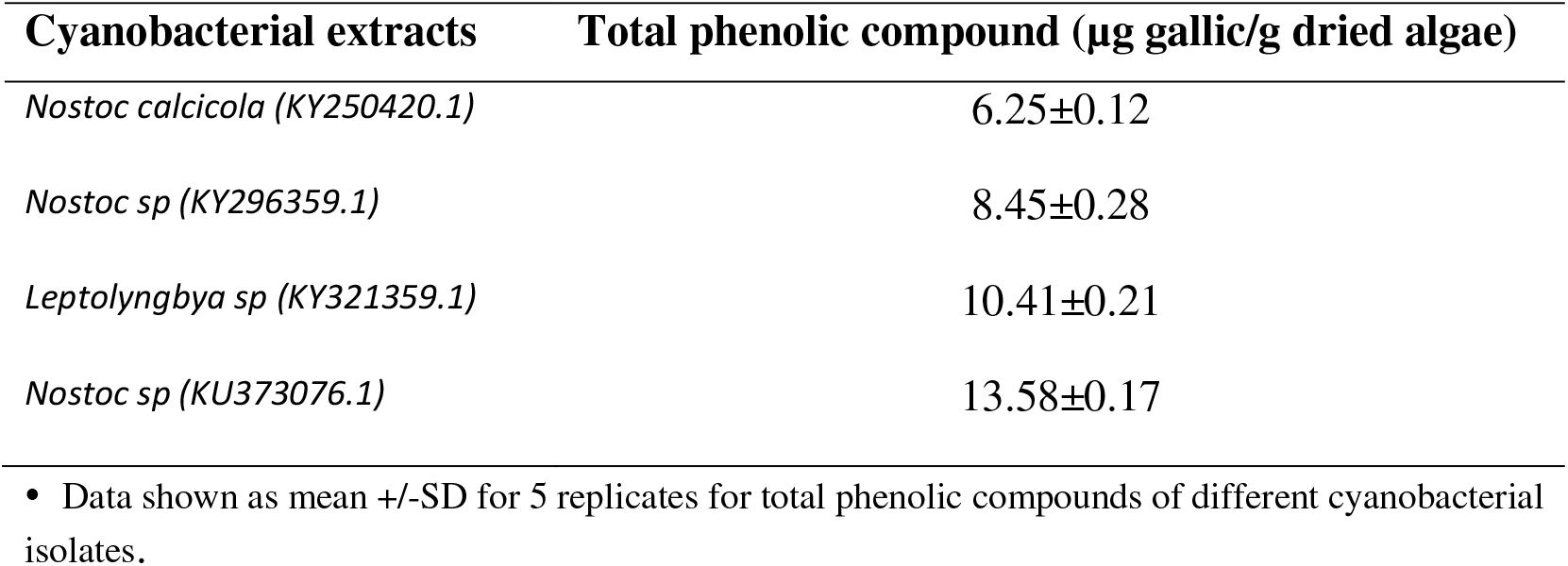
Total phenolic compounds in the extracts of obtained cyanobacterial isolates.

**Table 2.**
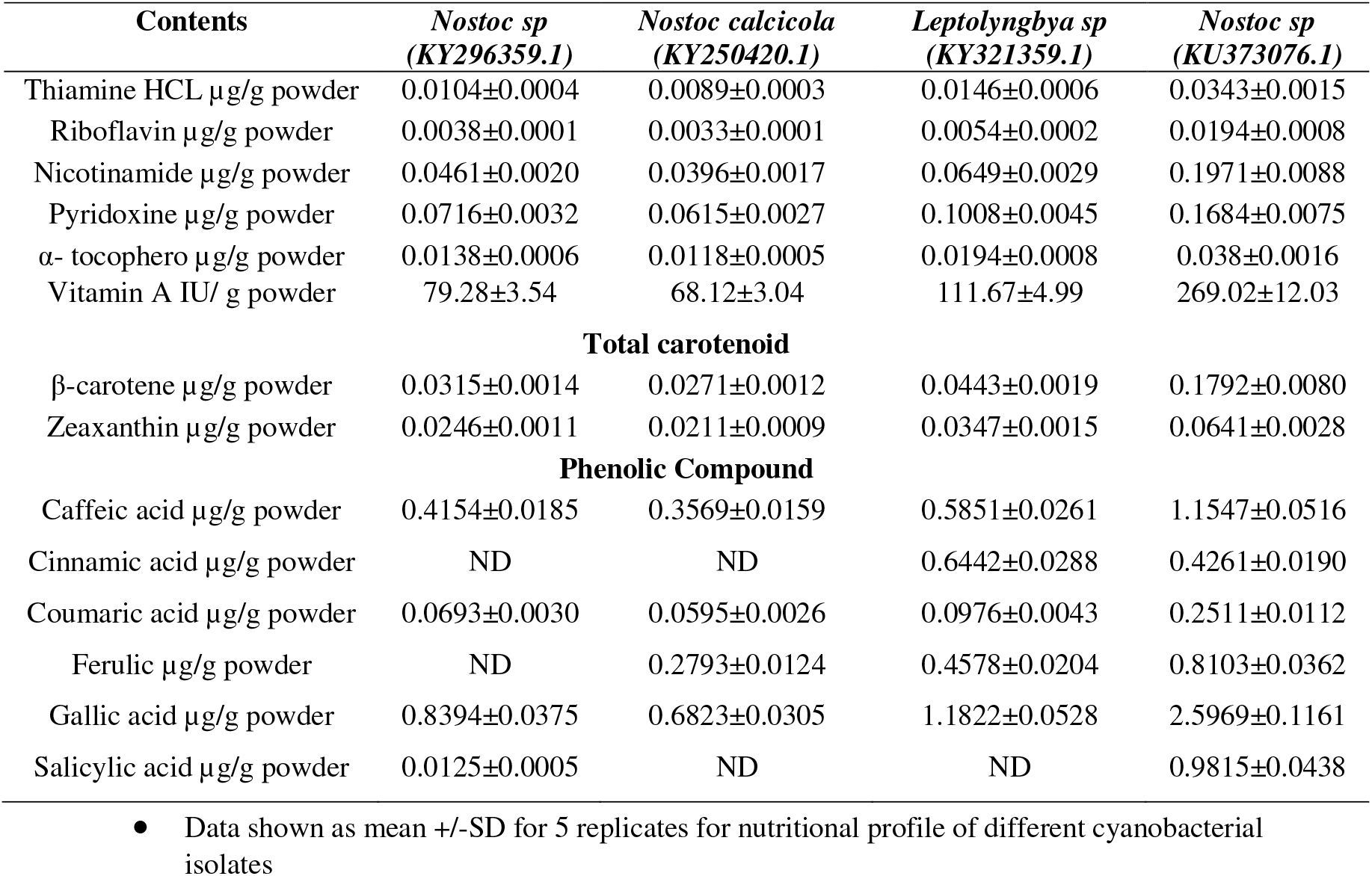
Nutritional profile of cyanobacterial isolates Powder

**Table 3.** DPPH radical scavenging activity (%) of cyanobacteria

| Cyanobacterial isolate | Concentration µg /ml | Scavenging activity % |
| --- | --- | --- |
| *Nostoc calcicola* (KY250420.1) | 50 | 18.25±0.37 |
| *Nostoc sp* (KY296359.1) | 50 | 19.16±0.39 |
| *Leptolyngbya sp* (KY321359.1) | 50 | 22.94±0.47 |
| *Nostoc sp* (KU373076.1) | 50 | 29.33±0.60 |

• Data shown as mean +/-SD for 5 replicates for scavenging activity of different Cyanobacterial isolates

### Wastewater treatment with the obtained cyanobacterial isolates

Cyanobacteria-based phycoremediation technologies have gained much attention recently as alternative bioremediation techniques over traditional methods for eco-friendly clean-up of metal-contaminated wastewater (Rawat et al., 2016) Cyanobacterial wastewater treatment is effective in the removal of nutrients (K, N and P) and heavy metals besides, the reduction of chemical and biological oxygen demand, removal of xenobiotic compounds and other contaminants (Olguín, 2003; Rawat et al., 2011; Abdel-Raouf et al., 2012; Cai et al., 2013). It is applicable to various types of wastewater including: human sewage, livestock wastes, agroindustrial wastes, industrial wastes, piggery effluent, food processing waste and other agricultural waste substrates (Abdel-Raouf et al. 2012; Cai et al. 2013). The produced wastewaters are increasing with the growing of human populations especially in developing countries, where the aquatic environments are suffering from contamination with heavy metals which is one of the most serious problems in Egypt (Zahran et al., 2015). Many previous studies stated the ability of different cyanobacterial species to treat waste water through removal of nitrogen, phosphors and heavy metals mainly by uptake these elements into algal cells as essential macronutrients for the growth of cyanobacteria (Aslan and Kapdan 2006; García et al., 2006; Ji et al., 2013). In this manner, the recent study of Foad et al. (2017) reported that dual bio treatment of *Chlorella vulgaris* and *Microccocus luteus* achieved the best removal of pollutants from wastewater via reduceing phosphorus by a percentage of (78.71%), nitrate (65.46%), potassium (49.9%) and magnesium (78.8%) within incubation period of 16 days. Also, Shanab et al. (2012) reported that *Pseudochlorococcum typicum* from aqueous solution showed percentage of metal bioremoval in the first 30 min of contact by 97% (Hg2+), 86% (Cd2+) and 70% (Pb2+). In this investigation the isolate *Nostoc sp* (KU373076.1) realized the best phycoremediation ability via reducing phosphorus, nitrate, potassium and magnesium by percentages of 96.7%,76.85%, 97.30% and 94.70%, respectively Table 4. Interestingly, all cyanobacterial isolates decreased lead, cadmium, Zinc and cobalt by 100 % (figure 7) and this may be due to collecting samples from Qalyubia irrigated drainage canals that receive contaminants from domestic, agricultural and industrial sources. According to Shanab et al. (2012) and Qari et al. (2014) algae appearing in polluted sites are either metal-tolerant or metal-resistant species. In the current study, the cyanobacterial isolates obtained from highly polluted sites with high concentration of heavy metals so, these isolates appeared high metal resistance with incomparable phycoremediation ability.

**Figure 7.**
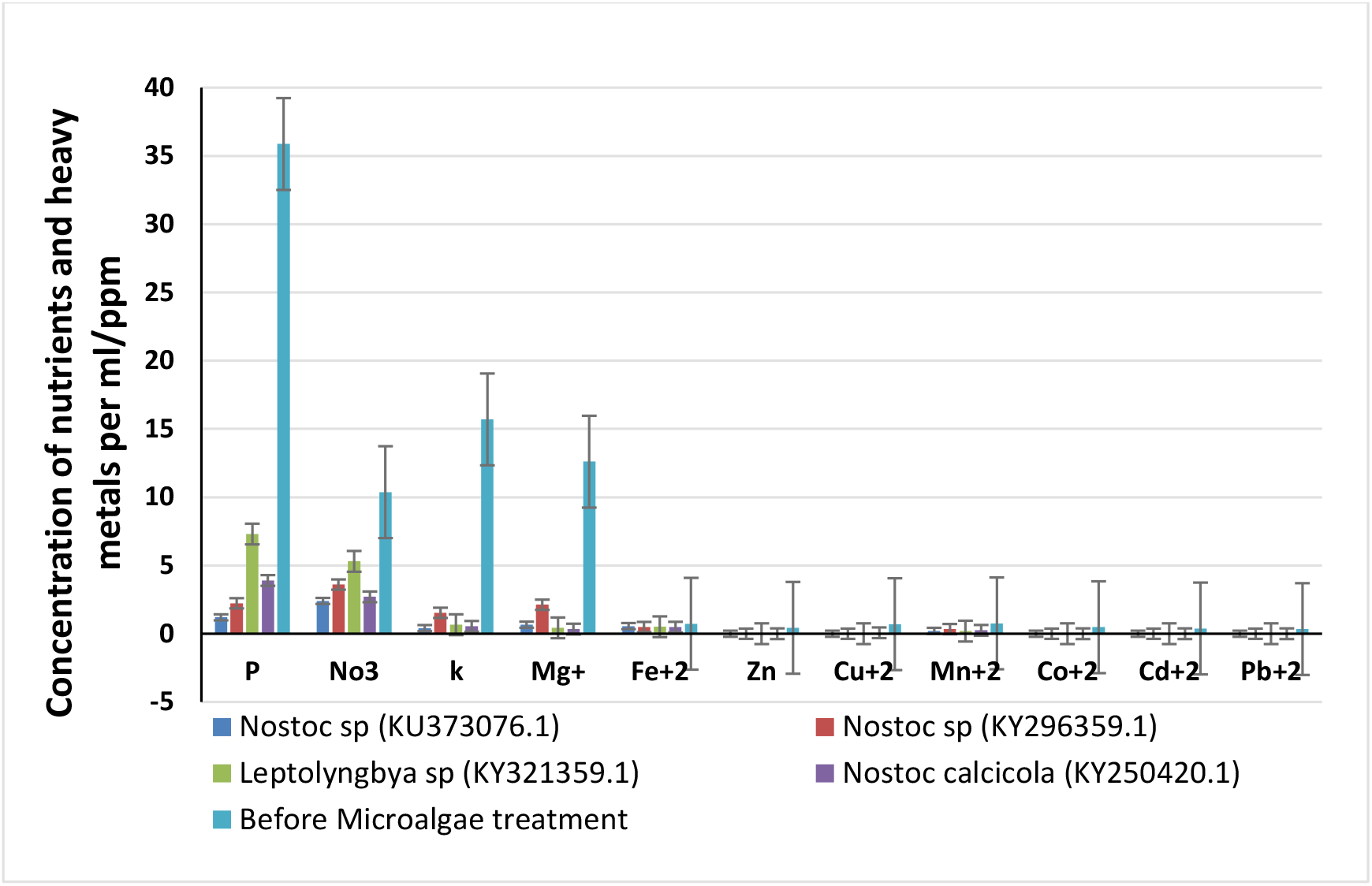
concentration of nutrients and heavy metals before and after cyanobacteria treatments.

**Table 4.** % of decreased heavy metals by the different cyanobacterial isolates

| **Metals**<br>**percentage** | *Nostoc calcicola*<br>**(KY250420.1)** | *Leptolyngbya sp*<br>**(KY321359.1)** | *Nostoc sp*<br>**(KY296359.1)** | *Nostoc sp*<br>**(KU373076.1)** |
| --- | --- | --- | --- | --- |
| P% | 89.2 | 80 | 93.8 | 96.7 |
| No3% | 73.9 | 49 | 65.3 | 76.85 |
| k% | 96.5 | 95.7 | 90.2 | 97.3 |
| Mg+% | 97.3 | 96.5 | 91.1 | 94.7 |
| Fe+2% | 34.2 | 30.1 | 32.8 | 23.2 |
| Zn% | 100 | 100 | 100 | 100 |
| Cu+2% | 90 | 100 | 100 | 100 |
| Mn+2% | 41 | 54.8 | 20 | 50 |
| Co+2% | 100 | 100 | 100 | 100 |
| Cd+2% | 100 | 100 | 100 | 100 |
| Pb+2% | 100 | 100 | 100 | 100 |

## Conclusion

The four identified cyanobacterial isolates exhibited antioxidant activity and Phycoremediation ability. The isolate *Leptolyngbya sp* could be novas species because of the low identity ratio with nearest sequences in database. On other hand, Nostoc *sp* (KU373076.1) realized the highest DPPH radical scavenging activity and this could be attributed to the presence of high phenolic constituents like caffeic and ferulic acids. All cyanobacterial isolates revealed incomparable phycoremediation ability since they obtained from stressed aquatic system where, there are different types of contaminates. Therefore, these cyanobacteria can be utilized either as agents for providing row martials for cosmeceuticals and pharmaceutical or as a promising cyanobacterial species for wastewater treatment. This study shed light on to what extent the environmental stresses could affect and alter cyanobacteria in aquatic systems.

## Acknowledgments

All thanks to Biotechnology Research Laboratory-2, Genetics and Genetic Engineering Department, Faculty of Agriculture, Benha University, www.bu.edu.eg.

